# Differential cellular stiffness contributes to tissue elongation on an expanding surface

**DOI:** 10.1101/2022.01.27.478117

**Authors:** Hiroshi Koyama, Makoto Suzuki, Naoko Yasue, Hiroshi Sasaki, Naoto Ueno, Toshihiko Fujimori

**Affiliations:** Division of Embryology, National Institute for Basic Biology (Div. Embryology, NIBB); SOKENDAI (The Graduate University for Advanced Studies); Division of Morphogenesis, National Institute for Basic Biology (Div. Morphogenesis, NIBB); Amphibian Research Center, Graduate School of Integrated Sciences for Life, Hiroshima University (ARC, Hiroshima Univ.); Laboratory for Embryogenesis, Graduate School of Frontier Biosciences, Osaka, University (FBS, Osaka Univ.)

**Keywords:** Pattern formation, morphogenesis, tissue elongation, cellular stiffness, vertex model, theory, mouse notochord

## Abstract

Pattern formation and morphogenesis of cell populations is essential for successful embryogenesis. Steinberg proposed the differential adhesion hypothesis (DAH), and differences in cell–cell adhesion and interfacial tension have proven to be critical for cell sorting. Standard theoretical models such as the vertex model consider not only cell–cell adhesion/tension but also area elasticity of apical cell surfaces and viscous friction forces. However, the potential contributions of the latter two parameters to pattern formation and morphogenesis remain to be determined. In this theoretical study, we analyzed the effect of both area elasticity and the coefficient of friction on pattern formation and morphogenesis. We assumed the presence of two cell populations, one population of which is surrounded by the other. Both populations were placed on the surface of a uniformly expanding environment analogous to growing embryos, in which friction forces are exerted between cell populations and their expanding environment. When the area elasticity or friction coefficient in the cell cluster was increased relative to that of the surrounding cell population, the cell cluster was elongated. In comparison with experimental observations, elongation of the notochord in mice is consistent with the hypothesis based on the difference in area elasticity but not the difference in friction coefficient. Because area elasticity is an index of cellular stiffness, we propose that differential cellular stiffness may contribute to tissue elongation within an expanding environment.

## Introduction

Pattern formation and morphogenesis by cell populations includes cell sorting, intermixing of different cell types, etc. These patterns are observed in various embryos and tissues such as germ layers, oviduct, and cochlea (1–4). A few hypotheses have been proposed to explain these phenomena, including the differential adhesion hypothesis (DAH) by Steinberg (5) in 1963 and differential interfacial tension hypothesis by Harris (6) in 1976. According to these hypotheses, either differential cell–cell adhesion or cell– cell interfacial tensions are considered, and their strengths are assumed to differ among cell types. These theories have been validated by both mathematical and experimental studies (4, 7, 8).

In mathematical studies, the vertex model and the Cellular Potts model are often used as a standard multicellular model, where interfacial tensions acting along cell–cell boundaries are considered a parameter (Fig. 1A, *λ*) (9, 10). Other basic parameters generally considered in these models are area elasticity of each cell and coefficient of viscous friction forces (Fig. 1A, *K*_a_ and *γ*, which will be introduced later) (9). The former denotes resistance against changes in apical cell surface area. For instance, when the apical cell surface is either stretched or compressed by external mechanical forces, the apical surface either increases or decreases, respectively, and the extent of the area change is determined by both the strength of the external forces and the area elasticity. Thus, this parameter is an index of cellular stiffness. On the other hand, the latter is derived from viscous friction forces exerted between cells and surrounding medium or tissues (11); increased friction forces restrict both cell movement and deformation. The friction forces between cells and surrounding tissues are affected by cell–cell interactions, cell– extracellular matrix interactions mediated by focal adhesions, etc (12–14). Although a spatial difference in friction forces is involved in the positioning cell populations in fish embryogenesis (12), the contributions of these two parameters to pattern formation and morphogenesis remain almost unknown.

**Figure 1:**
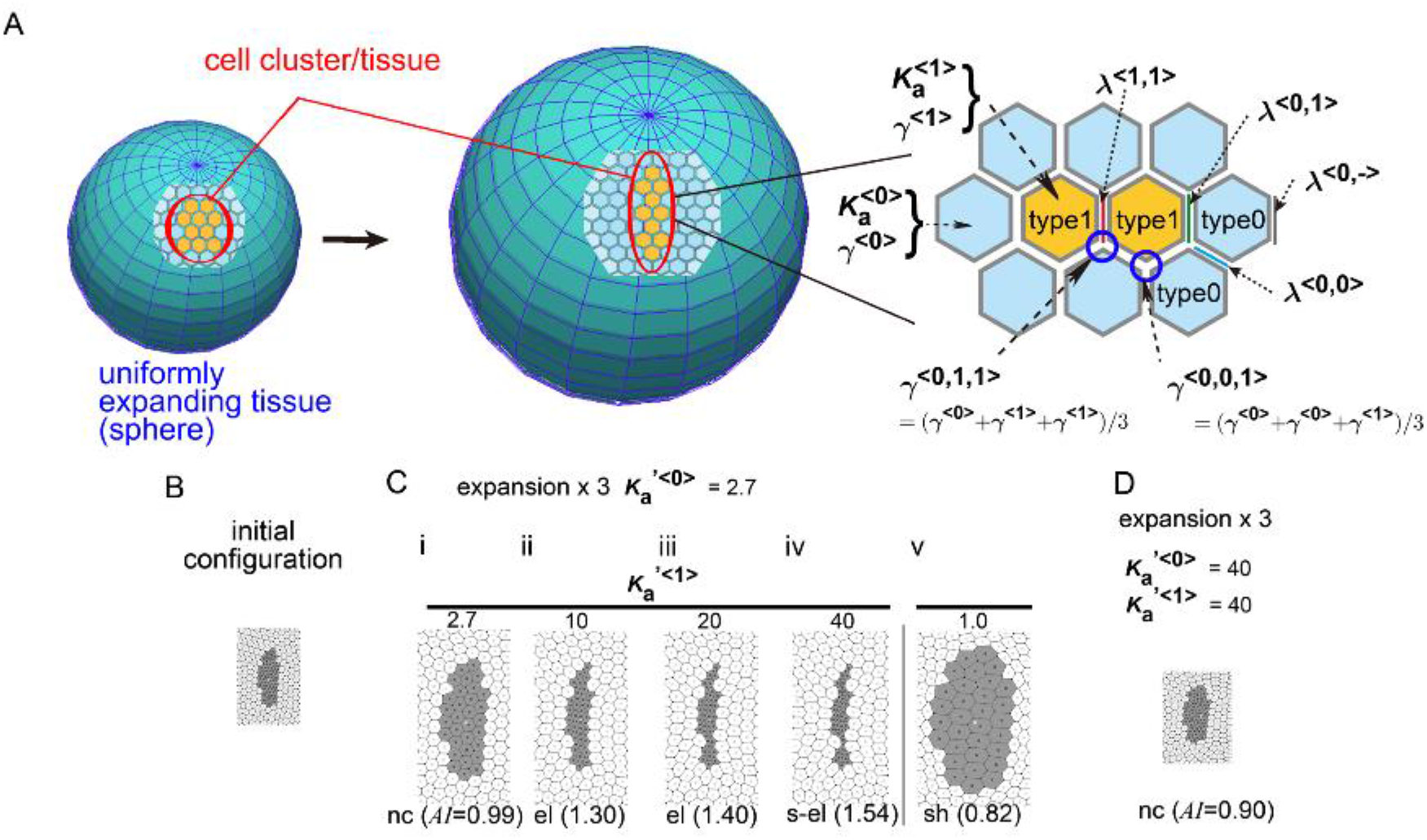
Difference in area elasticity causes elongation of cell cluster on expanding field. A. Two cell types with different area elasticity (*K*_a_) values were assumed in the vertex model; the area elasticity in type 1 cells (orange) and type 0 (light blue) cells are denoted as *K*_a_^<1>^ and *k*_a_^<0>^, respectively. These cells are placed on a uniformly expanding field (two spheres before and after expansion). Elongation of the type 1 cell cluster is the focus of this study. The line tensions between these cell types are shown (i.e., *λ*^<0,0>^, *λ*^<1,1>^, *λ*^<0,1>^, etc). The friction coefficients are shown for each vertex (blue circles; i.e., *γ*^<0,0,1>^, *γ*^<0,1,1>^), or for each cell (*γ*^<0>^ and *γ*^<1>^). B. The initial configuration of simulation is shown. The gray and white cells are type 1 and type 0, respectively. The whole xview of the configuration is provided in Fig. S1. C. Simulation outcomes under different values of 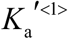 are shown (i–v; 2.7, 10, 20, 40, and 1.0). 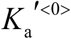 was fixed at 2.7. The outcome under the same value of *K*_a_’^<0>^ and *K*_a_’^1^ is also included (i; *K*_a_’^<0>^ = *K*_a_’^<1>^ = 2.7). The fields were expanded by three times in area. The indices of asymmetry/elongation (*AI*) relative to that of the initial configuration (B) are shown at the bottom of each panel (0.99, 1.30, 1.40, 1.54, and 0.82). According to these values, cell cluster shapes were classified as either “no change (nc)”, “elongated (el)”, “strongly elongated (s-el)”, or “shrunken (sh)” as defined in the main text. The scales of these images are the same as those in B. D. A simulation outcome achieved under high values of *K*_a_ s compared with C is shown (*K*_a_’^<0>^ = *K*_a_^<1>^ = 40). The fields were expanded by three times in area. The scale of the image is the same as that in B. Simulation results obtained using different area elasticity values are provided in Fig. S2.

We previously reported that in theory, on a uniformly expanding surface (Fig. 1A, an expanding sphere), a cell cluster is elongated even though directionally active cell movement is not assumed (Fig. 1A, a cell cluster in orange), whereas friction forces are considered between the cells and the expanding surface (15). This kind of expanding surface or field is analogous to an expanding rubber balloon in that the rubber membrane expands due to an increase in the volume of enclosed air, whereas a closed cell sheet expands by the increase in the volume of inner cavities such as amniotic cavities in mouse embryos (16). A cell sheet can also expand through proliferation of its component cells. On such an expanding field, even if the expansion is uniform or isotropic and a cell cluster has no intrinsic activity for directional migration, the cell cluster is elongated (15). We analyzed the mechanism with simulations and analytical approaches, as previously described (15). In our previous work, we assumed an isolated cell cluster, but, in typical epithelial tissues such as the mouse notochord, cells form a continuous cell sheet with other epithelial cell populations. For instance, the mouse notochord is surrounded by endodermal cell populations, resulting in a continuous cell sheet on the growing embryo. In the presence of such surrounding cell populations, we have not theoretically identified conditions for elongation of a cell cluster of interest.

In this study, we assumed a second cell population (Fig. 1A, blue cells) that surrounds a cell cluster of interest (Fig. 1A, orange cells), and determined theoretically if the cell cluster can be elongated. We found that the cell cluster can elongate, when either the area elasticity or the friction coefficient in the cell cluster is higher than that in the surrounding cell population. Moreover, we searched for *in vivo* phenomena where the above mechanism has been adopted. We previously found that elongation of the mouse notochord is promoted by expansion of the amniotic cavity (16). When comparing theoretical outcomes and experimental observations, the elongation of the mouse notochord is consistent with a difference in area elasticity, but not in friction coefficient.

## Theoretical model

### Vertex model

We adopted the simplest two-dimensional vertex model in which the mechanical potential energy (*U*) of a system is provided by line tensions of cell–cell interfaces and the area elasticity of each cell. A cell cluster of interest is defined as type 1 cells, and the surrounding cell populations as type 0 (Fig. 1A). The potential energy is defined as follows (15):

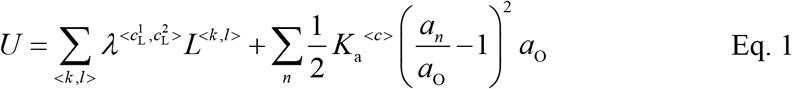

where *λ*^<*c^L^*_1_,*c^L^*_2_>^ and *L^<k,l>^* are the line tension and the length of the cell–cell interfaces between adjacent vertices *k* and *l* (Fig. 1), respectively. 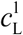 and 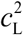 denote cell types of the two cells sharing the cell–cell interface <*k,l*> (Fig. 1A; *λ*^<1,1>^, *λ*^<0,1>^, etc.). *a_n_* is the area of the *n*th cell. *a_O_* and *K*_a_^<*c*>^ are a preferred area of the cell and the coefficient of area elasticity of type <*c*> cells, respectively (Fig. 1A; *K*_a_^<0>^ and *K*_a_^<1>^). The force (*F_h_*) exerted on an *h*th vertex is calculated as follows:

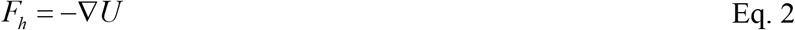

where ∇ is the nabla vector differential operator at each vertex. The motion of each vertex in polygonal cells is damped by friction forces and is described as follows:

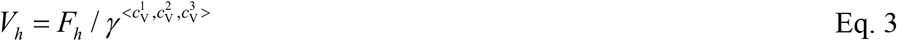

where *V_h_* is the velocity of the *h*th vertex, and 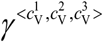 is the coefficient of the friction of a vertex that shares three cells whose types are denoted by 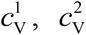, and 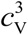 (Fig. 1A; *γ*^<0,0,1>^, *γ*^<0,1,1>^, etc.). The friction coefficients for each cell type are defined as *γ*^<0>^ and *γ*^<1>^ for type 0 and type 1 cells, respectively. The coefficient for each vertex was defined as the mean of the coefficients for the three cells: 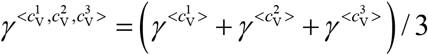(Fig. 1A). Okuda et al. (11) also assumed differential coefficients of friction. The friction is exerted between the cells and their surrounding medium or tissues. In this study, we assumed that the friction from the expanding field as defined below is dominant to that from the medium.

### A uniformly expanding field

A uniformly expanding field is analogous to the surface of an expanding rubber balloon; the area of the surface increases uniformly regardless of location on the surface. In the case of mouse early embryos around embryonic day 7.5, the volume of the inner cavity (e.g., the amniotic cavity) is increased, and thus the surface of the embryo is expanding. Cell proliferation within the surface tissue also causes it to expand (15). We adopted a simplifying assumption that a field expands in two dimensions. In this study, we placed a cell cluster on the expanding field with a surrounding cell population.

We previously defined the modeling of a uniformly expanding field (15). Briefly, when we arbitrarily defined a point on the two-dimensional field as an origin, other points are assumed to move away from the origin with speeds proportional to the distances between the points and the origin: *V_e_* ∝ *D_e_*, where *D_e_* is the distance between the point and the origin, and *V_e_* is the velocity. An object placed on this field also moves by *V_e_* if there are no other forces. Consequently, the equation that describes the motion of each vertex is modified as follows:

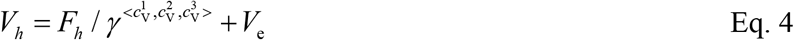

A similar formulation was previously proposed (11). We can interpret this equation as follows: 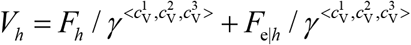, where *F_e|h_* is an apparent friction force provided by the expanding field. Under this assumption, we verified computationally and analytically that this expanding field does not yield any biased forces toward the cell cluster. Additionally, the expansion rate of the field was assumed to be temporally constant: *V*_e_ = *ε D_e_*, where *ε* is the expansion rate and is spatiotemporally constant. The parameters in our model were normalized by *λ*^<0,0^, 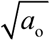, and *γ*^<0>^, and their dimensionless parameters are represented with a prime, e.g., 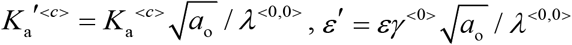.

## Results

### Differential area elasticity contributes to cell cluster elongation

We previously reported that a cell cluster (i.e., type 1 cells in this article) elongates on a uniformly expanding field in the absence of surrounding cell populations (the type 0 cells in this study). In real tissues, an epithelial cell cluster is not usually isolated but is instead surrounded by other cell populations. We theoretically searched for conditions under which the cell cluster can elongate even in the presence of surrounding cell populations.

First, we performed simulations using conditions under which type 1 and type 0 cells have the same parameter values, namely: *K*_a_^<0>^=*K*_a_^<1>^, *γ*^<0>^=*γ*^<1>^, and *λ*^<0,0>^ = *λ*^<1,1>^ = *λ*^<0,1>^. An initial configuration for the cells is shown in Fig. 1B, and the field was expanded. We set a slightly elongated initial configuration because we previously showed that this anisotropic configuration is a prerequisite for elongation (15). The cell cluster was enlarged due to friction forces from the expanding field (Fig. 1C-i). An index of asymmetry/elongation (*AI*) of a cell cluster was defined as described in the Materials and Methods section. If a cell cluster (type 1) is circular, *AI* becomes 1.0. If a cell cluster is elongated, *AI* becomes larger than 1.0. The *AI* relative to that of the initial configuration was 0.99 (Fig. 1C-i), indicating that the elongation of the cell cluster was not enhanced. We obtained a similar result with a different value for *K*_a_ s (Fig. 1D).

Next, we assigned different area elasticity values between type 1 and type 0 cells: *K*_a_^<0>^≠*K*_a_^<1>^ whereas *γ*^<0>^=*γ*^<1>^ and *λ*^<0,0>^ =*λ*^<1,1>^ If the area elasticity in type 1 cells was larger than that in type 0 cells (i.e., *K*_a_^1^ > *K*_a_^<0>^), elongation of the cell cluster was enhanced and the relative *AI*s were higher (Fig. 1C-ii, iii, and iv). In addition, if the area elasticity in the type 1 was smaller, the *AI* value of the cell cluster was decreased (Fig. 1C-v). These results indicate that, in theory, different area elasticity values contribute to tissue elongation.

We classified the morphological patterns of cell clusters by the value of the *AI* ratio after the simulation to its value before the simulation as follows: strong elongation (s-el; *AI_after_/AI_before_* > 1.5), elongation (el; *AI_after_/AI_before_* = 1.1-1.5), no change (nc; *AI_after_/AI_before_*= 0.9-1.1), and shrinkage (sh; *AI_after_/AI_before_* < 0.9). In Fig. 1C-D, these categories are written for each simulation outcome.

### Differential line tension between cell cell interfaces do not cause cell cluster elongation

We tested whether the DAH can reproduce the elongation of a cell cluster: *λ*^<0,0>^ ≠ *λ*^<1,1>^ ≠ *λ*^<0,1>^ whereas *K*_a_^<0>^ = *K*_a_^<1>^ and *γ*^<0>^ = *γ*^<1>^. We comprehensively examined the outcomes under different line tension values (Fig. 2A and 2C, the vertical and horizontal axes), and divided the parameter spaces between the morphological categories defined in Fig. 1. During classification, when more than two distinct cell clusters formed, the pattern was classified as either multi-cluster (cl; average cell number per cluster > 1.5) or intermixed (mx; average cell number per cluster < 1.5). As shown in Fig. 2A and 2C, we could not find conditions under which the cell cluster was elongated (Fig. 2); shrinkage occurred but not elongation of the cell cluster (Fig. 2, “sh”), and patterns with intermixing and multiple clusters were generated (Fig. 2, “mx” and “cl”, respectively). Thus, the differential line tensions between cell–cell interfaces alone do not cause tissue elongation.

**Figure 2:**
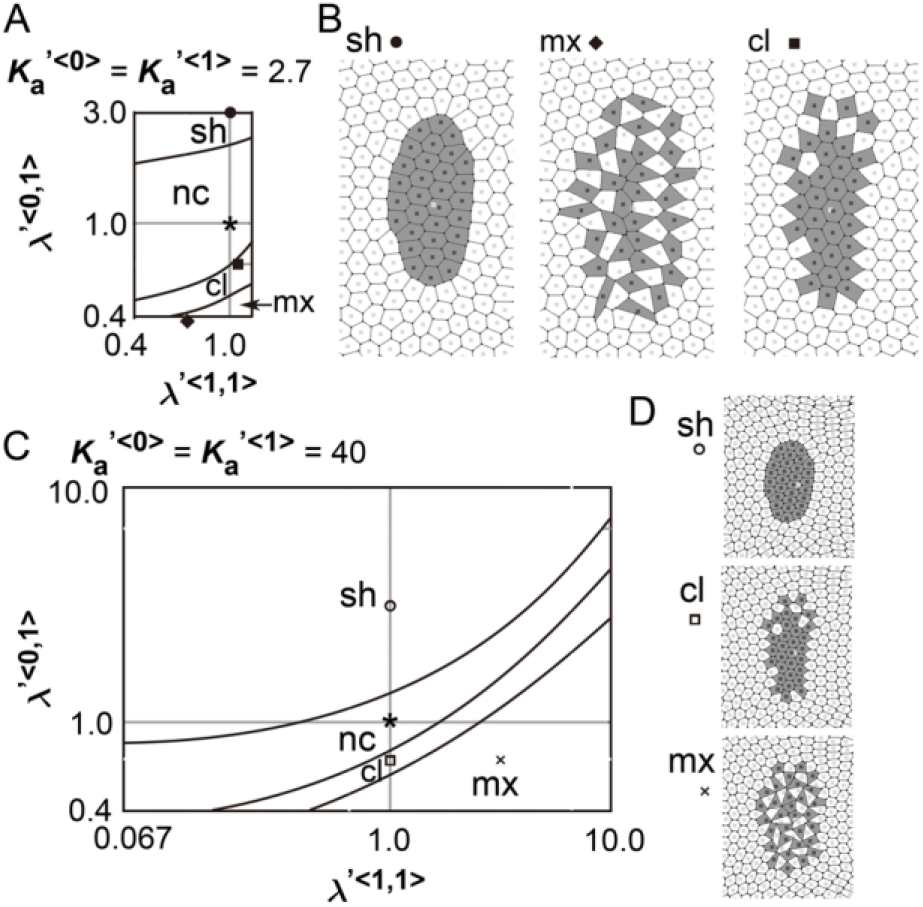
Difference in cell-cell adhesion does not cause elongation of cell cluster on expanding field. Simulations were performed using various line tension values, whereas the area elasticity and coefficient of friction were assigned the same values between the two cell types: *λ*^<0,0>^ ≠ *λ*^<1,1>^ ≠ *λ*^<0,1>^, whereas *K*_a_^<0>^ = *K*_a_^<1>^ and *γ*^<0>^ = *γ*^<1>^. The fields were expanded by three times in area. Simulation outcomes were shown in diagrams (A and C; the vertical and horizontal axes correspond to the line tensions). The parameter spaces in these diagrams were segmented according to the shapes of the cell clusters: shrinkage (“sh”), no change (“nc”), multi-cluster (“cl”), and intermixed (“mx”), which were defined in the main text. Some simulation outcomes are visualized in B and D, which correspond to A and C, respectively. The parameter values used for each condition in B and D are plotted on the diagrams (A and C), with symbols corresponding to each condition (circle, diamond, etc.). Simulation outcomes at the conditions with asterisks on the diagrams (A and C), where *λ*^<0,0>^=*λ*^<1,1>^=*λ*^<0,1>^, were previously shown in Fig. 1C-i and 1D, respectively. *K*_a_’^<0>^ = *K*_a_’^<1>^ = 2.7 (A); *K*_a_’^<0>^ = *K*_a_’^<1>^ = 40 (C).

### Optimal line tension between cell–cell interfaces are necessary in combination with area elasticity to induce cell cluster elongation

We analyzed the combinatorial effect of line tensions and area elasticity: *λ*^<0,0>^ ≠ *λ*^<1,1>^ ≠ *λ*^<0,1>^ and *K*_a_^<1>^ > *K*_a_^<0>^, whereas *γ*^<0>^ = *γ*^<1>^. Figure 3 shows the outcomes of simulations with different line tension values. The same condition as that in Fig. 1C-iv was plotted as an asterisk in Fig. 3A (right panel). The cell cluster was elongated and this elongation depended on field expansion (Fig. 3A, left panel vs. right panel). By changing the value of the line tension between type 1 and type 0 cells (i.e., *λ*^<0,1>^), the cell cluster either showed no elongation (“nc” and “sh”), formation of multiple clusters (“cl”), or intermixing (“mx”) (Fig. 3A, right panel). Similarly, by changing the value of the line tension between type 1 and type 1 cells (i.e., *λ*^<1,1>^), patterns other than elongation were generated. Typical configurations for each pattern (“sh”, “cl”, “mx”) are shown in Fig. 3B. These results indicate that, for the elongation induced by the differential area elasticity, optimal line tensions were required. Other outcomes under different conditions are shown in SI (Fig. S3 and S4).

**Figure 3:**
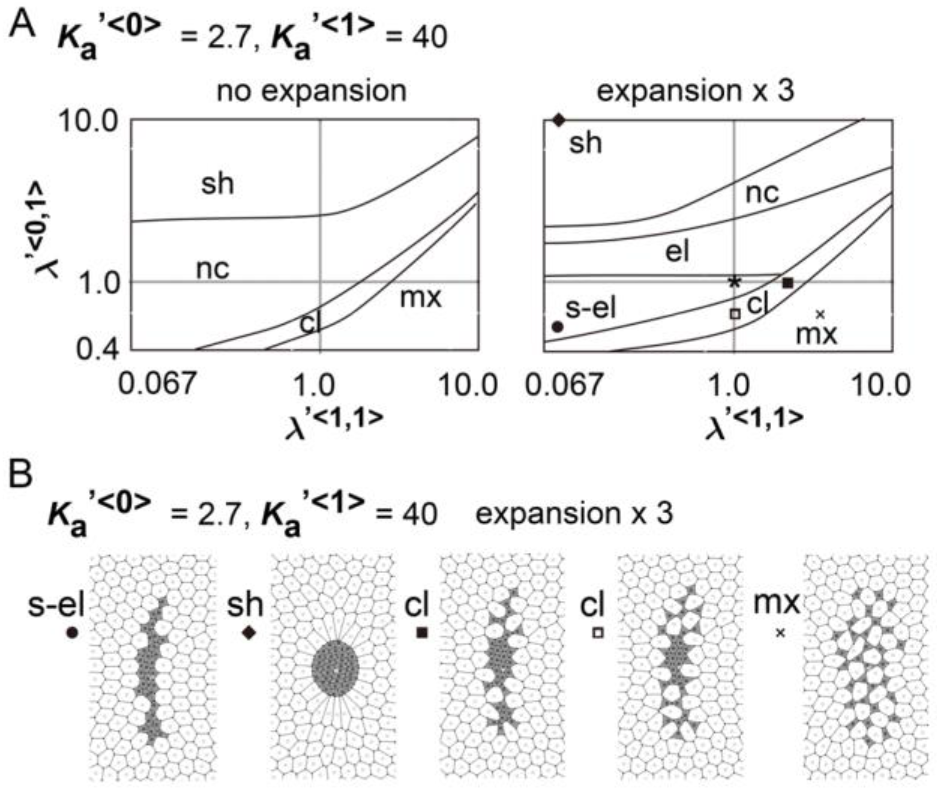
Difference in cell–cell adhesion contributes to cell cluster elongation with differential area elasticity on expanding field. Simulations were performed under conditions with various values for line tension and area elasticity, whereas the coefficient of friction was assigned the same value for both cell types: *λ*^<0,0>^ ≠ *λ*^<1,1>^ ≠ *λ*^<0,1>^ and *K*_a_^<0>^ ≠ *K*_a_^<1>^, whereas *γ*^<0>^ = *γ*^<1>^. Simulation outcomes are shown in diagrams in the absence and presence of the uniformly expanding field (left and right panels in A, respectively). In the panel on the right, the fields were expanded by three times in area. The vertical and horizontal axes correspond to line tensions, similarly to Fig. 2. The parameter spaces were segmented according to the shapes of the cell clusters similar to Fig. 2. The definition of the categories was described for Figs. 1 and 2 in the main text. Some simulation outcomes are visualized in B. The parameter values used for each condition in B are plotted on the diagrams (A), with symbols corresponding to each condition. *K*_a_’^<0>^ = 2.7; *K*_a_’^<1>^ = 40. Other simulation outcomes obtained with different values for the area elasticities are shown in Fig. S3. Simulation outcomes from a different initial configuration are shown in Fig. S4.

### Differential friction coefficient contributes to cell cluster elongation

We determined whether the differential coefficient of the friction forces causes a cell cluster to elongate: *γ*^<0>^ ≠ *γ*^<1>^, whereas *λ*^<0,0>^ = *λ*^<1,1>^ = *λ*^<0,1>^ and *K*_a_^<0>^ = *K*_a_^<1>^. The same condition as that in Fig. 1C-i was plotted as an asterisk in Fig. 4A; the cell cluster was not elongated. By increasing the value of the coefficient of friction among the type 1 cells (i.e., *γ*^<1>^), the cell cluster became elongated (Fig. 4A, “el”, and 4B, right-most panel marked with an open square). These results indicate that different values for the coefficient of friction contribute to tissue elongation.

**Figure 4:**
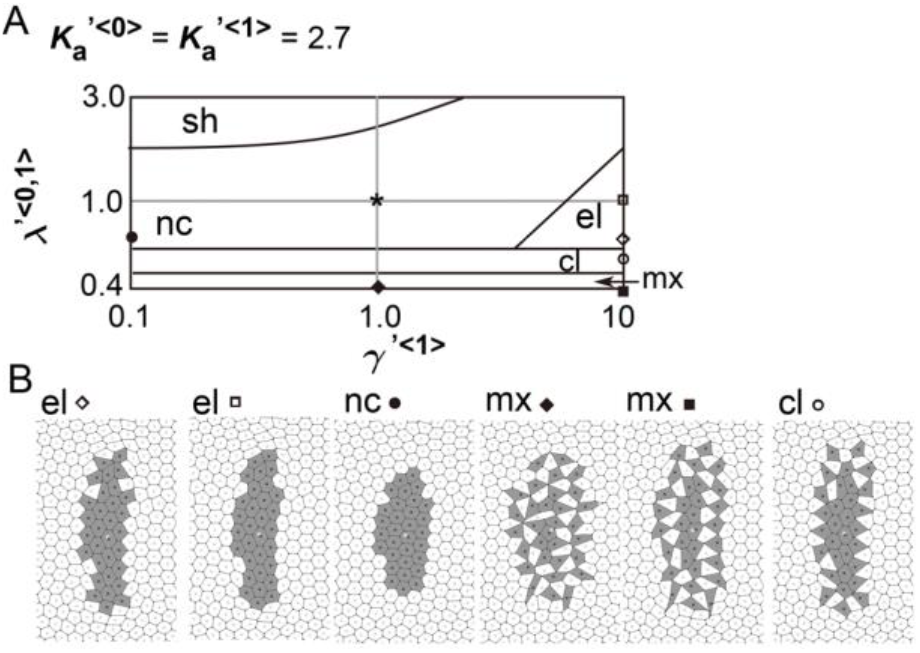
Differences in coefficient of friction cause cell cluster elongation on an expanding field. Simulations were performed under conditions with various values of the coefficient of the friction, whereas values for the area elasticity were assigned to be equal for both two cell types: *γ*^<0>^ ≠ *γ*^<1>^, whereas *K*_a_^<0>^ = *K*_a_^<1>^. The line tensions in the type 0 and type 1 cells were made equal in value, whereas the line tension between the type 0 and type 1 cells was different: *λ*^<0,0>^ = *λ*^<1,1>^ ≠ *λ*^<0,1>^. Simulation outcomes are shown in diagrams in the presence of the uniformly expanding field (A). The fields were expanded by three times in area. The vertical and horizontal axes correspond to the line tension and coefficient of friction, respectively. Parameter spaces were segmented according to the shapes of the cell clusters, similarly to Fig. 2, and the definition of the categories was described in the main text for Figs. 1 and 2. Some simulation outcomes are visualized in B. The parameter values used for each condition in B are plotted on the diagrams (A), with symbols corresponding to each condition. *K*_a_’^<0>^ = *K*_a_’^<1>^ = 2.7. A simulation outcome at the condition with an asterisk on the diagram (A) was previously shown in Fig. 1C-i. Other simulation outcomes obtained with different area elasticity values are shown in Fig. S5.

In addition, by changing the value of the line tension between the type 1 and type 0 cells (i.e., *λ*^<0,1>^), the cell cluster showed patterns of no elongation (“nc” and “sh”), multiple clusters (“cl”), and intermixing (“mx”) (Fig. 4A). Thus, optimal line tensions were required for the elongation induced by varying coefficients of friction force. Other outcomes under different conditions are shown in SI (Fig. S5).

### Cell behavior in mouse notochord elongation

In real tissues, elongation of various tissues is usually explained by directionally active cell movement that results in convergent extension (17, 18) where expanding fields are not considered. By contrast, we have shown experimentally that elongation of the mouse notochord depends on an increase in volume of the amniotic cavity (16). The mouse embryo on embryonic days 5.5–8.5 is either cylindrical or spherical in shape (Fig. 5A). The increase in the volume of this inner cavity pushes the surrounding cell layers that are composed of ectoderm, resulting in expansion of the ectodermal layer. This expansion is subsequently transduced to the outer cell layers, namely the mesoderm, endoderm, and notochord. The outermost layers in the mouse embryos are composed of the endoderm and notochord during the early stages of notochord formation (Fig. 5B). Therefore, the endoderm and the notochord would experience friction forces from expanding inner cell layers or the basement membranes between those layers.

**Figure 5:**
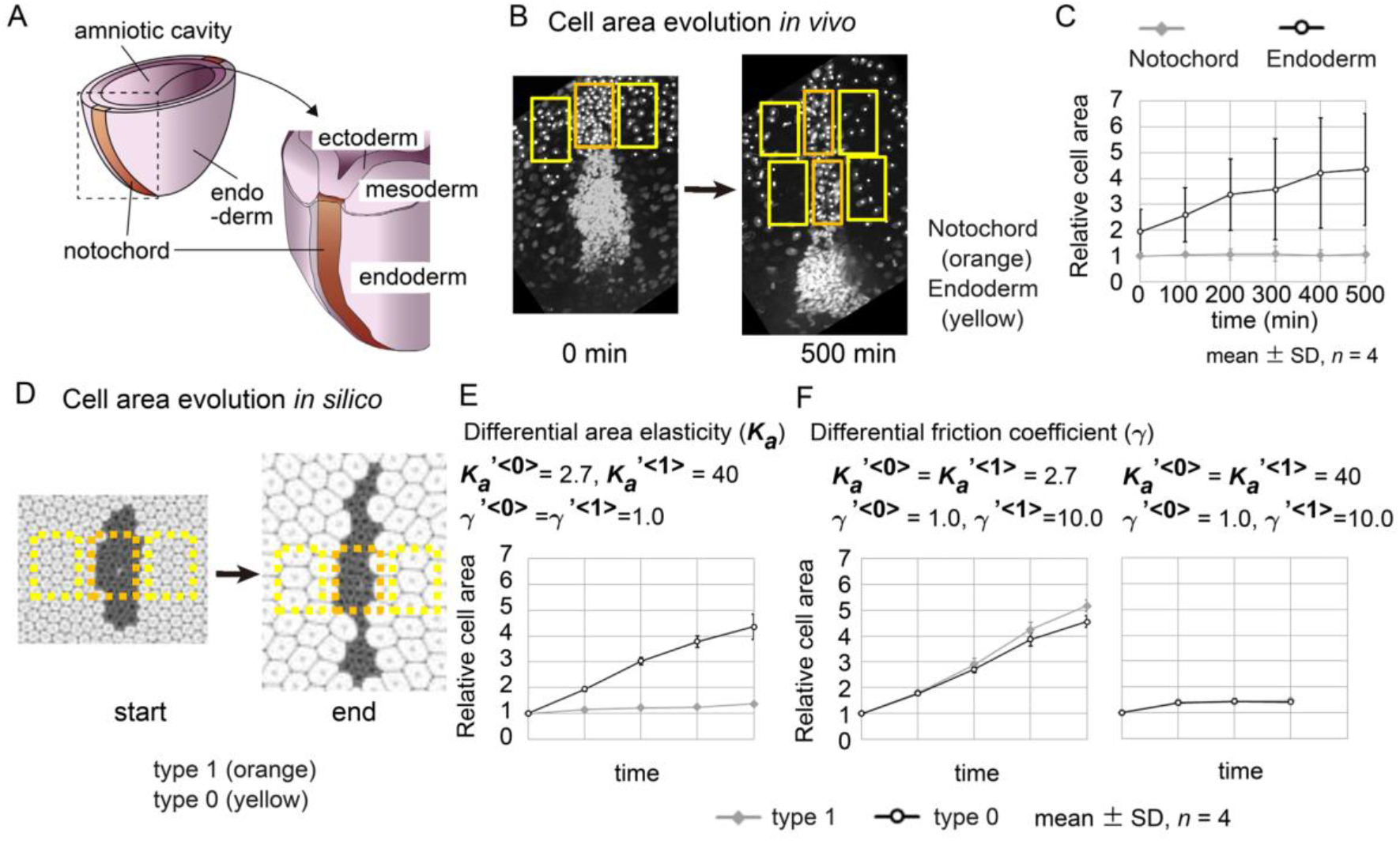
Cell behaviors in mouse notochord and their comparison with two theories A. Illustration of a mouse embryo with the notochord is shown. The embryo has an inner cavity (amniotic cavity) while the notochord and the endodermal layer form the outer surface of the embryo. Between the cavity and the outer surface, the ectodermal and mesodermal layers are found with basement membranes. This figure originated from our previous study (16). B. Confocal microscopic images at two time points in the mouse notochord and the surrounding endoderm are shown. Nuclei of the cells were visualized by histone 2B fused to EGFP. The orange and yellow rectangles are regions used for measuring cell areas within the notochord and endoderm, respectively. These microscopic images were obtained from our previous work (16). C. The cell areas in the notochord and endoderm are shown. The cell areas were estimated from the cell density in the rectangular regions in B, and the temporal evolutions were shown. The mean cell area in the notochord at 0 min was defined as 1.0. Four embryos were analyzed (*n* = 4). D. Cell areas in simulations were measured. Simulation outcomes are indicated with orange and yellow rectangles that were used to measure cell areas in a similar manner to B. E. Cell areas obtained with differential area elasticities, which were derived from the simulation outcomes in Fig. 1C-iv, are shown. Four different initial configurations of the simulations (*n* = 4) were applied. Parameter values for area elasticities and coefficients of friction are also indicated. F. Cell areas obtained with differential friction coefficients, which were derived from the simulation outcomes in Figs. 4 and S5, are shown in a similar manner to E.. The values from type 1 and type 0 cells are nearly identical.

From our theoretical analyses in Fig. 1 and 4, we raised two hypotheses for tissue elongation: area elasticity-based one, and friction coefficient-based one. We determined whether the elongation of the mouse notochord is consistent with differences in area elasticity or differences in friction coefficient. According to our previous data in Fig. 1C-iv and 4A, the cell area in the cell cluster of interest appears to either be almost unchanged according to the area elasticity–based hypothesis or increased according to the friction-based hypothesis. Conversely, the cell area in the surrounding cell populations seems to increase according to both hypotheses.

We went on to measure the dynamics of the cell area both in vivo and in silico. The apical cell area within the notochord was unchanged, whereas that in the endoderm increased (Fig. 5C). In the case of simulation data (Fig. 5D), based on the difference in area elasticity hypothesis, the cell area in the in silico cell cluster was unchanged, whereas that in the surrounding cell populations was increased (Fig. 5E). Based on the difference in friction coefficient hypothesis, the cell areas in both the cell cluster and the surrounding cell populations were larger (Fig. 5F). In addition, if the area elasticities in both cell types were made larger under different friction coefficients, cell area increases were restricted for both cell types but the dynamic was equivalent between the two cell types (Fig. 5F, a right panel). Thus, the dynamics of the cell areas according to the area elasticity–based hypothesis are consistent with the in vivo dynamics but not for the friction-based hypothesis.

### Experimental measurement of cellular stiffness in mouse notochord elongation

To further validate the area elasticity–based hypothesis in the mouse notochord, we measured cellular stiffness. To the best of our knowledge, no method for measuring area elasticity directly has been established to date. We used atomic force microscopy (AFM) that has been used to measure cellular stiffness (Young’s modulus) (19-21). A mouse embryo was placed on an agarose gel (Fig. 6B, light orange), and subsequently, a part of the embryo was overlaid by an additional agarose gel (Fig. 6B, dark orange). The Young’s modulus of the regions of the notochord or the surrounding endoderm was measured through indentation of the cantilever with a bead attached (Fig. 6C). A spatial map of the Young’s modulus was obtained (Fig. 6D). The Young’s modulus of the notochord regions was larger than that of the endodermal regions (Fig. 6E). These results suggest that the Young’s modulus of the notochord is higher than that of the endoderm.

**Figure 6:**
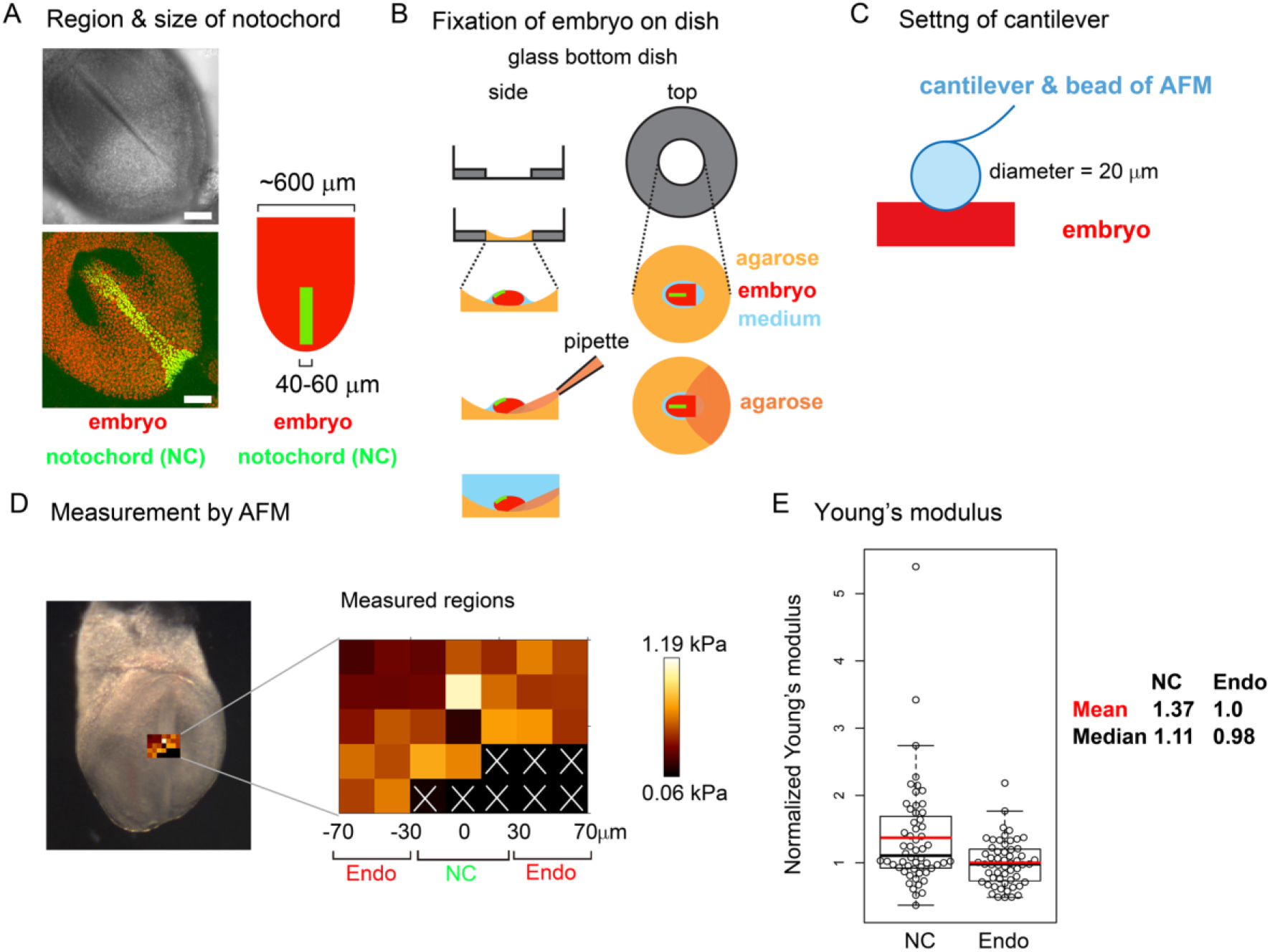
Young’s modulus measured using AFM in mouse notochord and endoderm A. Microscopic images of a mouse embryo are shown. Nuclei of the notochord cells were labeled with nuclear-EGFP (green), and nuclei of all embryonic cells including the endodermal cells as well as the notochord cells were labeled by histone 2B-mCherry (red). The surface of the mouse embryo is composed of the endodermal and notochord cells. Upper panel, brightfield; bottom panel, a maximum intensity projection image constructed from confocal fluorescent images; left panel, illustration of an embryo and the notochord. Typical widths of the notochord and embryo are written. B. Preparation procedure of embryo for AFM is illustrated. An embryo (red) is placed on an agarose gel (light orange) in a glass-bottom dish, and a part of the embryo is overlaid by an additional agarose gel (dark orange) before medium (blue) is added. Side and top views are shown. C. AFM cantilever assembly. A bead of 20 μm diameter was attached to the cantilever as described in the Materials and Methods section. D. A spatial map of Young’s modulus measured by AFM is shown. In the left panel, a brightfield microscopic image is provided where the regions subjected to the AFM measurement are also shown. In the middle panel, the Young’s modulus for each region in the embryo is shown with a 20 μm spatial interval. In regions marked by white crosses, AFM measurements failed to be carried out. The regions of the notochord and endoderm were estimated by the width of the notochord. NC, notochord; Endo, endoderm. In the right panel, a scale of Young’s modulus is shown. E. Young’s moduli in the notochord and the endodermal regions are compared. The mean value in the endodermal regions was set at 1.0, and the relative values were plotted. Four embryos with several data points were analyzed with total data points = 56 in both NC and Endo. The *p*-value calculated using the Mann–Whitney–Wilcoxon test was 0.006. Orange bars, mean; black bars, median; NC, notochord; Endo, endoderm.

## Discussion

In this study, we analyzed the elongation of a cell cluster on a uniformly expanding field using theoretical methods. In the case that the area elasticities or coefficients of friction differed between the cell cluster and the surrounding cell populations, the cell cluster was elongated as summarized in Table 1. By contrast, differences in cell–cell adhesion based on the differential adhesion hypothesis (DAH) cannot cause the cell cluster to elongate. The two hypotheses based on the area elasticity and the friction coefficient lead to different cellular behaviors; the apical cell areas in the surrounding cell populations are increased in both hypotheses, whereas the areas in the cell cluster of interest are either almost unchanged according to the former hypothesis or increased according to the latter (Table 1). Therefore, the elongation of the mouse notochord may be explained by the area elasticity–based hypothesis.

**Table 1:** Summary of comparison of phenomena; *in vivo* vs. different models on a uniformly expanding field

| Phenomena | *In vivo* | Uniformly expanding field |  |  |
| --- | --- | --- | --- | --- |
|  | mouse<br>notochord | Hypothesis A | Hypothesis B | Hypothesis C |
|  |  | Differential<br>cellular<br>stiffness | Differential<br>friction<br>coefficient | Differential<br>adhesion/tension |
| Elongation of cell<br>cluster/tissue | **+<sup>1</sup>** | **+** | **+** | **-** |
| Relative increase in cell<br>area (NC vs. END) | NC < END | NC < END | NC = END | not determined |

NC, notochord; END, endoderm
1. Imuta et al. (16)

Measurement of Young’s modulus through AFM suggests that the notochord is stiffer than the endoderm. The Young’s modulus of the notochord and endoderm differed by ~1.4 times, whereas the difference in the area elasticity in our simulations was up to 10 times. The Young’s modulus differs from the area elasticity, although both are measures of cellular stiffness. During the AFM-based measurement, the direction of indentation is parallel to the apico–basal axis. On the other hand, the area elasticity is related to the change in apical cell area whose direction is perpendicular to the apico– basal axis. Nevertheless, the change in apical cell area and in the apico–basal height should be related under conserved cell volume; the increase of apical cell area should lead to the decrease in the apico–basal height, and vice versa. Although we do not have quantitative relationship between the Young’s modulus and the area elasticity, we suppose that the Young’s modulus reflects, at least partially, the area elasticity. In general, cell stiffness can differ by over an order of magnitude (22, 23). The notochord in chordates is believed to provide stiffness of their bodies (24, 25), and the notochord in *Xenopus laevis* was experimentally shown to be several to several tens times stiffer than the endoderm (23).

Mechanisms of tissue elongation have been experimentally and theoretically studied well (17, 18). In these mechanisms, a cell cluster of interest is assumed to have an intrinsic activity of directional cell movement or anisotropic property of cell–cell interfaces (10, 17, 26), whereas any expanding field is not considered. Our present study shed light on a possible contribution of an expanding field to pattern formation, and consequently, the involvement of the area elasticity and the coefficient of the friction in tissue elongation was demonstrated. Expansion of a cavity and its role in morphogenesis has been discussed in both mouse and fish (27, 28). Friction between fields and cells should exist in development of various multicellular systems including germ layers (12, 29, 30), epidermis during pregnancy (31), and cells in contact with other cell layers such as smooth muscle layers or with external structures such as eggshells (14, 32, 33). Cell-extracellular matrix interaction is important for morphogenesis (34, 35) and would also be related to the friction forces. Further investigation is required to clarify what kind of real tissues our two hypotheses apply to.

## Materials & methods

### Mathematical model and analysis

The implementation of our mathematical model is essentially the same as that in our previous article. The surrounding cell populations have an outer boundary as shown in Fig. S1, and cropped views are shown in Figs. 1–4. Total simulation time 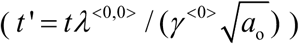 was fixed at 30 as a dimensionless time. Simulations were performed using the Euler method.

The definition and measurement of the asymmetry/elongation index (*AI*) were reported previously (15). The length of the longest axis of a cell cluster was measured as the maximum caliper. *AI* was defined as *AI* = *Feret* / *D*_circle_, where *Feret* is the maximum caliper, and *D*_circle_ is the diameter of a circle with the same area as the cell cluster. Thus, *AI* is 1.0 in a circle and is increased in an elongated shape. Simulation outcomes were converted to TIFF images, and the *Feret* and the area of a cell cluster were measured using ImageJ (*Feret* is prepared as a measurement option for ImageJ. https://imagej.nih.gov/ij/docs/menus/analyze.html#set).

The definition of the patterns in Figs. 1–4 is described in the main text. Briefly, when all type 1 cells formed a single cluster, the pattern was categorized as either “s-el”, “el”, “nc”, or “sh”, according to the ratio of *AI* after the simulation to *AI* before the simulation; when more than two separate clusters formed, the pattern was categorized into either “cl” or “mx”.

### Image analysis for estimating cell area

Cell areas in the mouse notochord and endoderm were estimated as follows. A rectangular region was defined on the notochord at 0 min (Fig. 5B, left panel, 0 min, orange rectangle). Rectangular regions with the same width as the above were defined on the endodermal regions, which were also adjacent to the rectangle on the notochord (Fig. 5B, 0 min, yellow rectangles). For images after time evolution (Fig. 5B, 500 min), rectangular regions were defined on the notochord, whose widths may differ from that at 0 min. Rectangular regions set on the endodermal regions at 500 min have the same widths as at 0 min. Cell areas in these regions were defined as [the area of the rectangle / the nuclear density]. Similar procedures were also carried out for simulation outcomes as shown in Fig. 5D with orange and yellow rectangles.

### Mouse embryo

The notochord cells were identified by the expression of *Brachyury* (*T*). The *Brachyury*-expressing cells were labeled by nuclear enhanced green fluorescent protein (EGFP) as reported previously (36) (Acc. No. CDB0604K; http://www2.clst.riken.jp/arg/mutant%20mice%20list.html). Briefly, knock-in mice expressing both Brachyury and nuclear EGFP from the endogenous *Brachyury* gene locus were used. All embryonic cells including the endodermal cells were labeled with mCherry-fused H2B (histone 2B proteins) expressed under the control of a ubiquitous promoter, *ROSA26*, as we reported previously (37). By mating these two mouse lines with mice from a subsequent generation, we eventually obtained a mouse line that is both homozygous for *H2B-mCherry* and heterozygous for *Brachyury* with the nuclear *EGFP* gene. By mating males from this mouse line with ICR female mice (Japan SLC), embryos expressing both H2B-mCherry and nuclear EGFP were obtained with 50% probability.

### Atomic force microscopy (AFM)

Embryos described above were isolated on embryonic day 7.5. The embryos were placed in DMEM containing HEPES with 50% FBS on ice. The concentration of the agarose (BMA, SeaKem GTG, cat. 50070) in Fig. 6B is 1% melted in PBS. The embryos were placed on the agarose (Fig. 6B, light orange), and the medium was almost removed. Finally, a small amount of 1% agarose was added to anchor the embryos (Fig. 6B, dark orange). DMEM containing HEPES with 50% FBS was added on ice again. Thirty minutes before the AFM measurement, the above embryos were transferred to an incubator at 37°C, and the DMEM medium was replaced with PBS just before the AFM measurement. Four distinct embryos were subjected to AFM measurement, and for each embryo, several tens of measurement points were defined as described in Fig. 6D. Data points that did not yield a clear force-indentation curve were omitted from the data analysis (Fig. 6D, white crosses).

We could not identify the exact location of the notochord during AFM, because our AFM has a bright field microscope but not a good fluorescent one. Alternatively, we independently performed a fluorescent imaging using a confocal fluorescent microscope (Nikon A1, Japan) as shown in Fig. 6A, and estimated the width of the notochord at 40-60 γm. Because the midline of the notochord was distinguishable in the bright field microscopy of the AFM (Fig. 6D), we assumed that the notochords were located around the midline and 40-60 μm width.

AFM measurements were conducted as previously described (20). In brief, a JPK Cellhesion 200 (Bruker) fitted with an x/y-motorized stage and mounted on a macro zoom microscope (Axio Zoom.V16, Zeiss) was used. Customized AFM probes (Novascan) were prepared by attaching borosilicate beads (Fig. 6C, 20 μm diameter) to tipless rectangular silicon cantilevers (350 μm long, 32.5 μm wide, 1 μm thick; nominal spring constant 0.03 N/m, MikroMasch). Force-distance curves (maximum indentation force: 3 nN, approach speed: 5 μm/s) were acquired every 20 μm apart in a bidirectional raster scan (Fig. 6D), leading to that data points on the three columns adjacent to the midline were expected to be on the notochord. Cell elasticity (Young’s modulus) values on the tissue surface were calculated based on the Hertz model and mapped onto brightfield images using the JPK data processing software (Bruker).

## Author contributions

H.K. designed the work; H.K., H.S, and T.F. contributed to the conception; H.K. designed models; H.K. developed computational algorithms; H.K. prepared embryos for AFM measurements; M.S., N.Y., and N.U. designed and performed AFM experiments; H.S. provided mouse live imaging data; all authors wrote the manuscript; all authors contributed to the interpretation of the results.

## Acknowledgements and Funding

This work was supported by KAKENHI from the Japan Society for the Promotion of Science (JSPS) for T.F. and H.K., and the NINS program for cross-disciplinary science study for H.K.

