## Supplementary information for "Differential cellular stiffness contributes to tissue elongation on an expanding surface"

**Koyama et al.**

This file includes Supplementary Figures S1–S5.

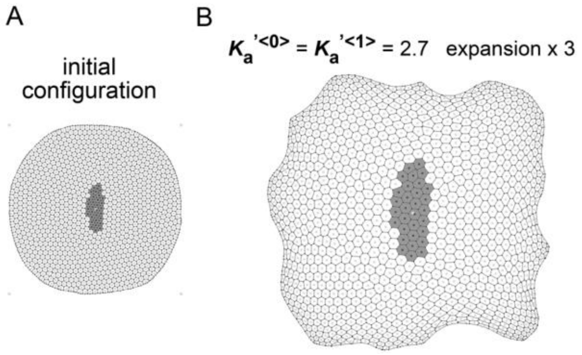


Figure S1 (related to Fig. 1): Overview of simulation.

A. Overview of the initial configuration of simulations in Figs. 1–4.

B. Overview of the simulation outcome in Fig. 1C-i.

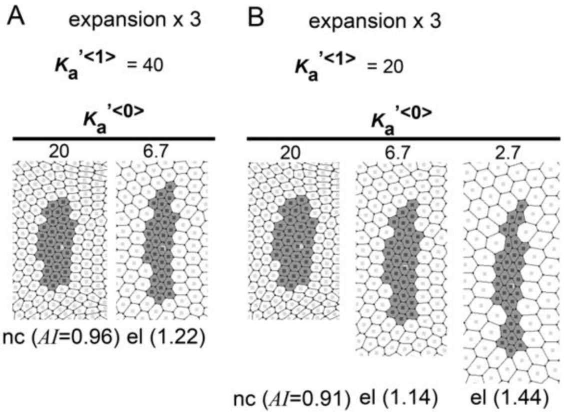


Figure S2 (related to Fig. 1): Simulation outcomes given various area elasticity values.

A. Simulation outcomes given different area elasticity values for type 0 cells ( $K_a{}'^{<0>}$ ) (20 and 6.7). The area elasticity in the type 1 cells ( $K_a{}'^{<1>}$ ) was set at 40. The fields were expanded by three times in area. The indices of asymmetry/elongation *(AI*) relative to that of the initial configuration (Fig. 1B) are shown at the bottom of each panel in a similar manner to Fig. 1C. These images are all of the same scale.

B. Simulation outcomes under different values of $K_a{}'^{<0>}$ are shown (20, 6.7, and 2.7). $K_a{}'^{<1>}$ was set at 20. The fields were expanded by three times in area. The indices of asymmetry/elongation *(AI*) are shown in a similar manner to A. These images are all of the same scale.

In both A and B, the cell cluster was only elongated under conditions with $K_a{}^{<1>} > K_a{}^{<0>}$ but not with $K_a{}^{<1>} < K_a{}^{<0>}$.

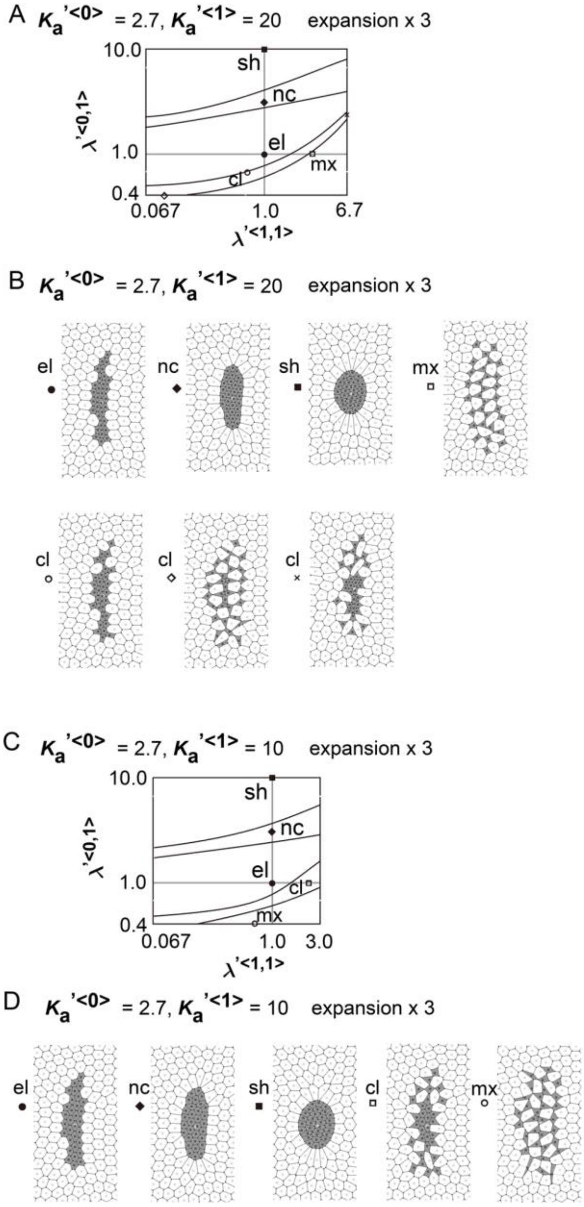


Figure S3 (related to Figure 3): Effect of differences in cell–cell adhesion on elongation of cell cluster given various values for area elasticities.

Simulations were performed in a similar manner to Fig. 3, except for different combinations of the values of the area elasticities in the type 1 and 0 cells. In A and B, $K_a'^{<1>}$ was set at 20, whereas, in C and D, $K_a'^{<1>}$ was set at 10. The phase diagrams (A and C) showed an almost similar pattern to Fig. 3A.

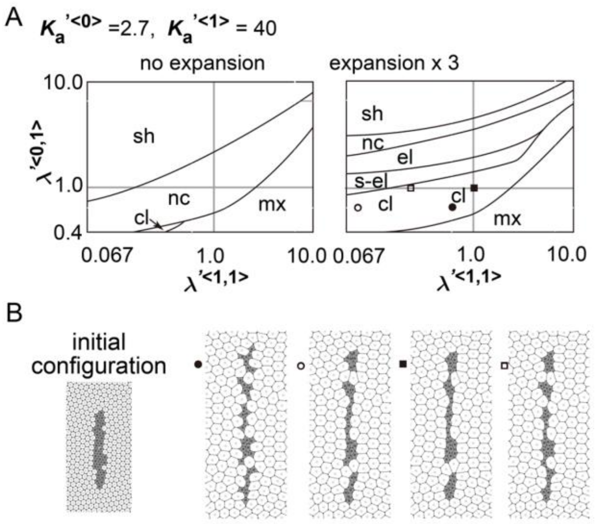


Figure S4 (related to Fig. 3): Difference in cell–cell adhesion contributes to pattern formation of cell cluster with differential area elasticity on an expanding field.

Simulations were performed in a similar manner to Fig. 3, except for a different initial configuration of simulations (B, initial configuration), which forms a more elongated shape compared with that in Fig. 1B. The phase diagrams (A) show a nearly identical pattern to Fig. 3A. In addition, as shown in B, a single cluster of cells sometimes formed a linear array of multiple distinct cell clusters, and such a pattern is observed in various developing tissues (1, 2).

1. S. M. Meilhac, *et al.*, A retrospective clonal analysis of the myocardium reveals two phases of clonal growth in the developing mouse heart. *Development* **130**, 3877–3889 (2003).
2. E. Tzouanacou, A. Wegener, F. J. Wymeersch, V. Wilson, J.-F. Nicolas, Redefining the Progression of Lineage Segregations during Mammalian Embryogenesis by Clonal Analysis. *Dev. Cell* **17**, 365–376 (2009).

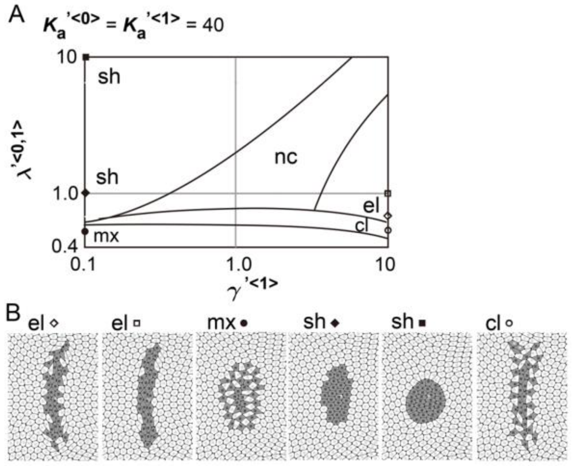


Figure S5 (related to Fig. 4): Difference in coefficient of friction causes cell cluster elongation on an expanding field.

Simulations were performed in a similar manner to Fig. 4, except for different area elasticity values:

$K_a'^{<0>} = K_a'^{<1>} = 40$. The phase diagram (A) shows a nearly identical pattern to Fig. 4A.
